# Components of the endocytic and recycling trafficking pathways interfere with the integrity of the *Legionella*-containing vacuole

**DOI:** 10.1101/781849

**Authors:** Ila S. Anand, Won Young Choi, Ralph R. Isberg

## Abstract

*Legionella pneumophila* requires the Dot/Icm translocation system to replicate in a vacuolar compartment within host cells. Strains lacking the translocated substrate SdhA form a permeable vacuole during residence in the host cell, exposing bacteria to the host cytoplasm. In primary macrophages, mutants are defective for intracellular growth, with a pyroptotic cell death response mounted due to bacterial exposure to the cytosol. To understand how SdhA maintains vacuole integrity during intracellular growth, we performed high-throughput RNAi screens against host membrane trafficking genes to identify factors that antagonize vacuole integrity in the absence of SdhA. Depletion of host proteins involved in endocytic uptake and recycling resulted in enhanced intracellular growth and lower levels of permeable vacuoles surrounding the Δ*sdhA* mutant. Of interest were three different Rab GTPases involved in these processes: Rab11b, Rab8b and Rab5 isoforms, that when depleted resulted in enhanced vacuole integrity surrounding the *sdhA* mutant. Proteins regulated by these Rabs are responsible for interfering with proper vacuole membrane maintenance, as depletion of the downstream effectors EEA1, Rab11FIP1, or VAMP3 rescued vacuole integrity and intracellular growth of the *sdhA* mutant. To test the model that specific vesicular components associated with these effectors could act to destabilize the replication vacuole, EEA1 and Rab11FIP1 showed enhanced colocalization with the vacuole surrounding the *sdhA* mutant compared with the WT vacuole. Depletion of Rab5 isoforms or Rab11b reduced this aberrant colocalization. These findings are consistent with SdhA interfering with both endocytic and recycling membrane trafficking events that act to destabilize vacuole integrity during infection.

## Introduction

*Legionella pneumophila* is a fastidious Gram-negative bacterium that grows intracellularly in both fresh water amoebae and human alveolar macrophages (Isberg *et al*., 2009). Disease in humans is initiated after inhalation of aerosolized water particles contaminated with the bacterium (Cunha *et al*., 2016). Upon entry into the lungs, *L. pneumophila* is internalized by alveolar macrophages with its most severe manifestation being Legionnaires’ disease pneumonia (Horwitz & Maxfield, 1984; Horwitz & Silverstein, 1980). In both amoebae and macrophages, the ability of *L. pneumophila* to replicate intracellularly is dependent on the Icm/Dot type IV secretion system (T4SS) (Segal *et al*., 2005). This complex introduces more than 300 translocated substrates into the host cell cytosol, which promote establishment of the *Legionella*-containing vacuole (LCV) and exert a variety of regulatory controls on the host cell (Asrat *et al*., 2014; Burstein *et al*., 2016).

Establishment and maintenance of LCV membrane integrity is essential for protection of *L. pneumophila* from innate immune cytosolic sensing (Creasey & Isberg, 2014; Liu *et al*., 2018). The inability of a bacterial mutant to properly form the LCV or maintain its integrity results in a severe intracellular growth defect and pathogen clearance (Creasey & Isberg, 2012; Wiater *et al*., 1998). The identification of the Icm/Dot-translocated substrate SdhA has greatly contributed to our understanding of the consequences of failure to properly maintain pathogen replication vacuoles (Laguna *et al*., 2006). In the absence of SdhA, the LCV becomes disrupted and *L. pneumophila* is detected in the cytoplasm of the infected macrophage (Creasey & Isberg, 2012; Laguna *et al*., 2006). Bacteria exposed in this fashion to the cytosol are susceptible to recruitment of the interferon-regulated GBP family of anti-microbial proteins, resulting in leakage of LPS and activation of caspase-11 (Creasey & Isberg, 2012; Liu *et al*., 2018; Piro *et al*.). As a consequence, gasdermin D is activated and pyroptotic cell death ensues (Pilla *et al.,* 2014; Aachoui *et al*., 2014; Shi *et al*., 2015).

We previously demonstrated that mutations that removed the bacterial phospholipase PlaA rescued the defect in vacuole integrity observed with the D*sdhA* mutant, resulting in increased numbers of intracellular bacteria, thus demonstrating that PlaA promotes vacuole disruption (Creasey & Isberg, 2012). PlaA bears homology to the translocated phospholipase SseJ from *Salmonella* typhimurium, which is also involved in vacuole disruption in the absence of another *Salmonella* translocated substrate, SifA (Ruiz-Albert *et al*., 2002; Akoh *et al*., 2004; Flieger *et al*., 2002; Ruiz-Albert *et al*., 2002). During infection, SifA binds host factor SKIP and sequesters Rab9 to prevent delivery of M6PR cargo to the vacuole (McGourty *et al*., 2012). The resemblance between SdhA/PlaA and SifA/SseJ suggests that SdhA may regulate host membrane trafficking events to maintain LCV integrity.

In addition to *Salmonella*, other pathogens have been reported to interface with host membrane trafficking pathway as a strategy for maintaining vacuolar integrity. The *Chlamydia* protein IncD was shown to recruit the host CERT-VAP complex to the inclusion and enable acquisition of host lipids that are essential for inclusion membrane stability (Derre *et al*., 2011). Recently, the *Shigella* vacuole has been reported to interact with endocytic Rab5-positive vesicles and recycling Rab11-positive vesicles to facilitate vacuole membrane rupture, indicating that vacuole integrity can be disrupted by interfacing with appropriate cellular compartments (Mellouk *et al*., 2014; Weiner *et al*., 2016). In this study we determine if the disruption of LCV integrity that results from loss of *L. pneumophila* SdhA function can be attributed to a subset of host membrane trafficking pathways. Using high throughput screening strategies, we identified specific endocytic and recycling Rab GTPase isoforms and their downstream effectors as playing a role in vacuole disruption resulting from challenge with the Δ*sdhA* strain. The similarity between *Legionella* and *Shigella* interactions with a common endocytic-recycling pathway provides an example of how a host process can either promote or interfere with pathogen replication, depending on the strategy used for intracellular growth.

## Results

### Identification of host proteins that contribute to disruption of *L. pneumophila* Δ*sdhA* intracellular growth

To identify host membrane trafficking pathways that interfere with growth of the *L. pneumophila ΔsdhA* mutant, we carried out a screen using an siRNA library targeting genes known to participate in membrane trafficking and remodeling. RAW 264.7 macrophages are known to be defective in the activation of pyroptosis and clearance in response to cytoplasmic exposure of *L. pneumophila* pattern recognition molecules, so growth of the mutant could be followed without premature host cell death downstream of cytoplasmic bacterial exposure (Pelegrin *et al*., 2008). This allowed subtle differences in intracellular growth to be detected. RAW 264.7 macrophages were seeded in 96-well plates and transfected with an arrayed siRNA library against 112 mouse genes involved in membrane trafficking (**Supplemental Tables 1 and 2**). Each well contained 4-pooled siRNAs per gene target and each plate contained 6 non-targeting siRNA control wells. After 48 hours of transfection, cells were challenged with a Δ*sdhA* Lux^+^ strain and luminescence was measured as a proxy for intracellular growth, performing the assay in at least triplicate samples for each gene target (**Figure 1A**). Candidates were identified by comparing the luminescence from each siRNA-treated well to the average luminescence of the non-targeting siRNA controls. The median absolute deviation score (ZMAD) was then determined for each gene target. We focused our analysis on early time points 12hpi and 24hpi, becaise siRNA knockdown effects diminish after 72 hours of transfection, to identify candidates that stimulate Δ*sdhA* intracellular growth after depletion.

**Figure 1.**
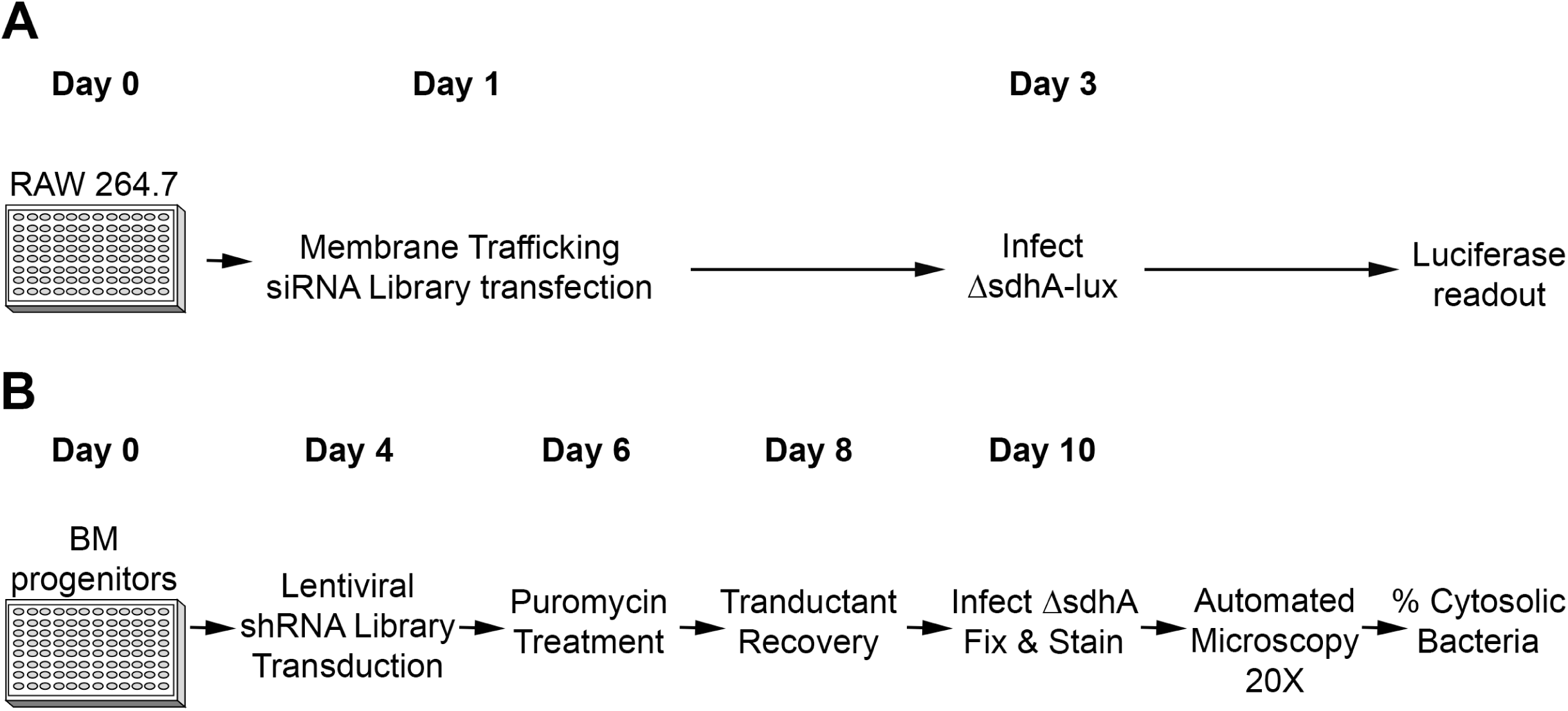
Loss of function treatments that compensate for depressed vacuole integrity of intracellular *L. pneumophila* Δ*sdhA.* (A) Identification of siRNA that enhance intracellular growth. RAW 264.7 macrophages were seeded in 96-well plates and transfected the following day with an siRNA library directed against transcripts encoding membrane trafficking proteins. After 2 days treatment, transfected cells were challenged with *L. pneumophila* Δ*sdhA* Lux^+^ and luminescence was measured at 12 and 24 hours post infection (hpi). (B) Identification of shRNA that decrease vacuole permeability in primary macrophages. A/J bone-marrow derived progenitors were seeded in 96-well plates and incubated in differentiation medium for 4 days. Cells were transduced with shRNAs targeting genes identified in the siRNA screen (A). Transductants were selected by puromycin treatment for 2 days and recovered by further incubation in puromycin-free medium. Terminally differentiated BMDM transductants were challenged with Δ*sdhA* for 6h. Plates were fixed and stained for cytosol-detected *L.p.* and total *L.p.*. High content imaging was performed with 20X objective and percent of cytosol-detected bacteria for each well was measured by image analysis.

A total of 20 genes were identified as hits in the siRNA screen based on increased intracellular growth relative to the nontargeting siRNA controls with a cutoff of ZMAD ≥ 1.5. Candidate genes spanned functional roles in endocytosis, endocytic recycling, and exocytosis (**Figure 2A**). Hits included arrestin subtypes (ARRB1 & ARRB2), clathrin heavy chain (CLTC), regulators of actin cytoskeleton (ROCK1, CDC42, DNM2, ARFIP2, WASF1, VAV2), E3 ubiquitin ligase (NEDD4L), phosphoinositide kinase (PI3K), components of vesicle fusion (VAMP1 & SYT1), and specific Rab GTPase isoforms (RAB3D, RAB4B, RAB5B, RAB5C, RAB8A, RAB11B). Interestingly, only five genes were identified as strong hits at both 12hpi and 24hpi (ARRB1, RAB3D, RAB4B, RAB5C, SYT1). Hits identified at 12hpi may alter pathways that are toxic for growth of bacteria during early steps of vacuole biogenesis, while hits identified at later times (such as DNM2 encoding a dynamin isoform) could block pathways that cause decay to an already established vacuole.

**Figure 2.**
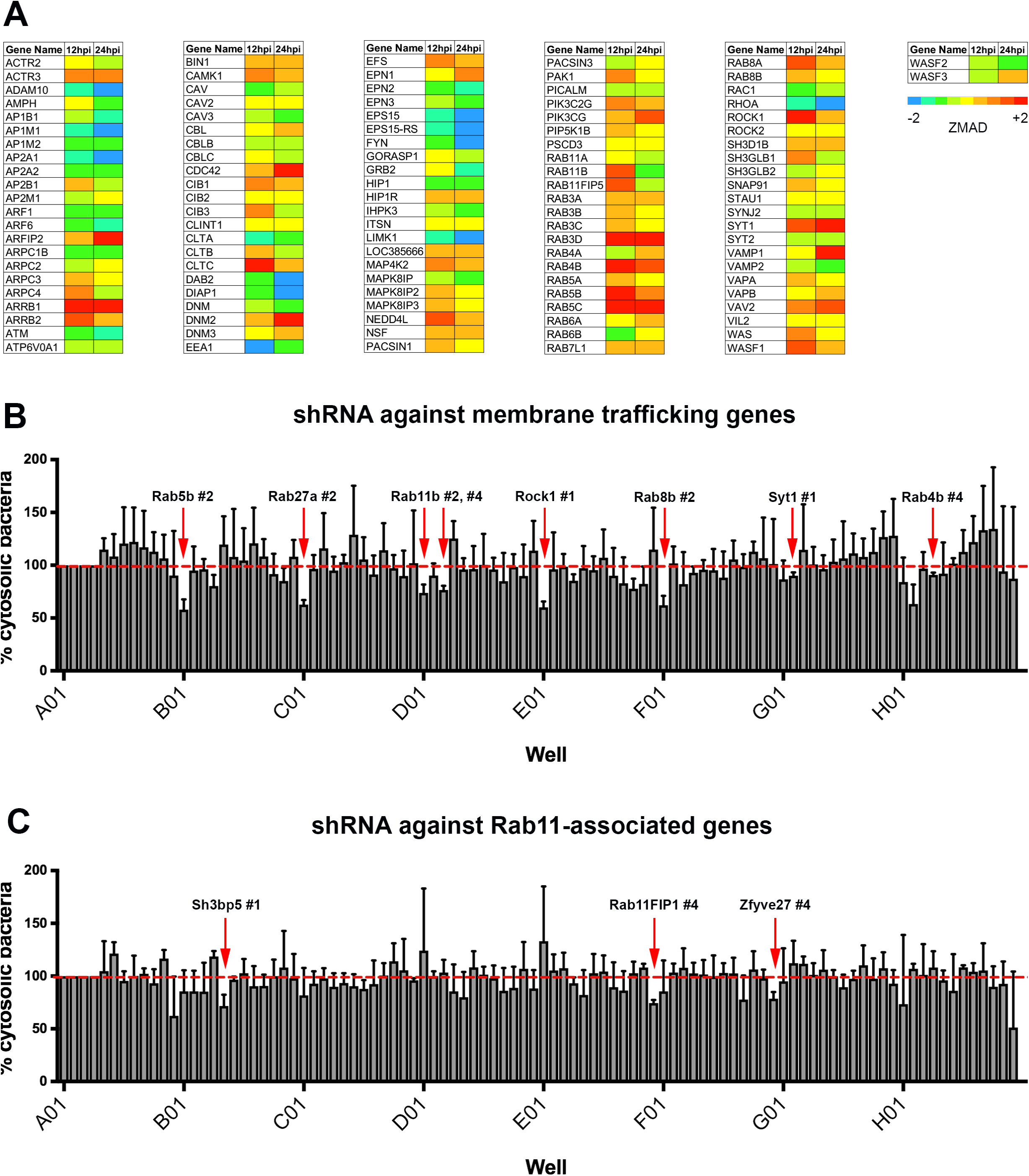
RNAi screens identify membrane trafficking proteins that antagonize *L. pneumophila* Δ*sdhA* intracellular growth and vacuole integrity. (A) Results of high-throughput siRNA screen for enhanced intracellular growth of *L. pneumophila* Δ*sdhA* Lux^+^. Experimental design, Fig 1(A), performing luminescence readings at 12 or 24 hpi, with least three replicate measurements for each gene target being performed Luminescence from each experimental well was normalized to the average luminescence of the non-targeting siRNA treated control wells of the same plate. Normalized ZMAD values are shown color-coded for each siRNA of the library. A cutoff of ZMAD ≥ 1.5 was used to select potential candidates. (**B)** Candidates identified in 2(A) and additional targets not covered in the original screen were tested in a high-throughput shRNA screen, performing assay detailed in Fig. 1B, detecting cytosolic exposure of bacteria. Three replicates for each knockdown condition were performed, capturing 16 images/well with automated microscopy. Image capture, analysis and normalization of data were performed as described (Experimental Procedures). **(C).** Depletion of transcripts for a subset of Rab11 effectors results in increased vacuole integrity during *L. pneumophila* Δ*sdhA* intracellular growth. Assay conditions as in panel 2B. Statistical analyses were performed on normalized data by unpaired t test (*P < 0.05) (Experimental Procedures).

### Identification of shRNAs that partially rescue Δ*sdhA* vacuole integrity

The assay for increased bacterial yields allowed identification of knockdown candidates that could improve intracellular growth of the Δ*sdhA* mutant, but they may not increase the frequency of intact vacuoles. To directly test if improved growth was due to increased vacuole integrity, we selected hits from the siRNA screen and additional related targets for analysis in a secondary screen using a direct assay for vacuole integrity in bone marrow-derived macrophages (BMDM) from the mouse. The secondary screen took advantage of a fluorescence readout we previously described to detect disrupted Δ*sdhA* vacuoles in knockdown macrophages combined with shRNA knockdown in these primary cells (Creasey & Isberg, 2012). Terminally differentiated BMDMs were used to facilitate microscopic readout, and to avoid the complication of having a portion of the cells in S phase, which results in a high proportion of unstable vacuoles (de Jesus-Diaz *et al*., 2017). This strategy allowed subtle differences in vacuole integrity to be detected. BMDM were transduced with a lentiviral shRNA library (**Figure 1B**) targeting 23 membrane trafficking genes arrayed as 4 individual shRNAs assayed separately (**Supplemental Table 2**). Lentivirus encoding sh-LacZ was used as a control. After allowing differentiation and 6 days knockdown, BMDMs were challenged in triplicate with the Δ*sdhA* strain for 6 hours, fixed and probed for permeable vacuoles using the antibody accessibility assay (Creasey & Isberg, 2012). After capturing multiple images per well and subjecting them to image analysis to quantitate permeable vacuoles, shRNAs that increased the integrity of the Δ*sdhA* vacuole relative to the shRNA-LacZ negative control were identified (**Figure 2B**).

In this fashion, eight shRNAs were identified that significantly reduced the number of permeable Δ*sdhA* vacuoles detected in macrophages (Table 1, t test, P = 0.0048-0.0346). The candidate genes included five members of the Rab family of GTPases that are associated with endocytic recycling, as well as ROCK1, and SYT1. Of this set, Rab5b, Rab11b, and Rab8b were of particular interest due to their robust phenotype and their tight association with recycling, based on STRING analysis (Franceschini *et al*., 2013). Strikingly, two hits from the screen were shRNAs directed against Rab11b (Table 1), so this result was pursued further. Using the same screening procedures as described above, we performed a lentiviral shRNA library screen against gene targets related to Rab11 (**Figure 2C**). These gene targets included Rab11 guanine exchange factors (SH3BP5 & TRAPPC2L), motor proteins (MYO5B, KIF5A, KIF5B, KIF13A), adaptor protein (ZFYVE27), components of the exocyst complex (EXOC1-8), and downstream Rab11-family interacting proteins (RAB11FIP1-5) (**Supplemental Table 3**). ShRNAs against SH3BP5, ZFYVE27, and RAB11FIP1 significantly reduced the number of disrupted Δ*sdhA* vacuoles detected in macrophages (Table 1; P<0.0041 for RAB11FIP1). The gene target Rab11FIP1 was of particular interest, since this downstream Rab11 effector is known to regulate membrane trafficking events and vesicle docking events (Horgan & McCaffrey, 2009; Welz *et al*., 2014). We selected Rab5b, Rab11b, Rab8b, and Rab11FIP1 for further characterization in response to a Δ*sdhA* infection.

**Table 1.** Hits from shRNA lentiviral screens

| Figure | Well | shRNA | Description/Function | P value |
| --- | --- | --- | --- | --- |
| 2B | B01 | Rab5b #2 | Rab-GTPase. Endocytosis | 0.0172 |
| 2B | C01 | Rab27a #2 | Rab-GTPase. Exocytosis | 0.0048 |
| 2B | D01 | Rab11b #2 | Rab-GTPase. Endocytic Vesicle Recycling | 0.0293 |
| 2B | D03 | Rab11b #4 | Rab-GTPase. Endocytic Vesicle Recycling | 0.0091 |
| 2B | E01 | Rock1 #1 | Rho-associated Protein Kinase | 0.006 |
| 2B | F01 | Rab8b #2 | Rab-GTPase. Endocytic Vesicle Recycling | 0.0181 |
| 2B | G02 | Syt1 #1 | Synaptotagmin 1 | 0.0346 |
| 2B | H04 | Rab4b #4 | Rab-GTPase. Endocytic Vesicle Recycling | 0.0172 |
| 2C | B05 | Sh3bp5 #1 | Rab11 GEF. Endocytic Vesicle Recycling | 0.044 |
| 2C | E12 | Rab11FIP1 #4 | Rab11 Effector. Endocytic Vesicle Recycling | 0.0041 |
| 2C | F12 | Zfyve27 #4 | Protrudin. Endocytic Vesicle Recycling | 0.281 |

### Interfering with early endosomal traffic stabilizes the LCV harboring the Δ*sdhA* mutant

Host membrane trafficking events are coordinated by small Rab GTPases in the cell (Pfeffer & Aivazian, 2004). Rab5 is involved in regulating endocytic trafficking events and Rab11 and Rab8 regulate recycling events and anterograde transit (Stenmark, 2009). Little is known about how the *Legionella* vacuole subverts interaction with these Rabs and, specifically, their individual isoforms. A role for Rab5 isoforms in destabilizing the LCV, in particular, was surprising because previous work with this protein favored a model in which Rab5 acts to interfere with intracellular growth by driving the LCV into an endocytic or degradative compartment (Gaspar & Machner, 2014; Ku *et al*., 2012; Lucas *et al*., 2014). To determine if reduced Rab5 function can stabilize the vacuole harboring the D*sdhA* strain, BMDMs were nucleofected with siRNAs against individual Rab isoforms and knockdown efficiency was determined by immunoblot (**Figure 3**). Individual or group depletion of Rab5a, Rab5b, or Rab5c resulted in enhanced vacuole integrity in BMDM challenged with *L. pneumophila* Δ*sdhA* (**Figure 3A**). Furthermore, each individual knockdown resulted in enhanced intracellular growth of the Δ*sdhA* strain. In particular, initiation of intracellular replication was clearly revived in a subset of cells treated with the siRNA, although it was clear that growth restoration was far from complete (**Figure 3B**). This result was consistent with our previous experiments showing that increasing vacuole integrity in the absence of SdhA function only allows partial restoration of intracellular growth (Creasey & Isberg, 2012).

**Figure 3.**
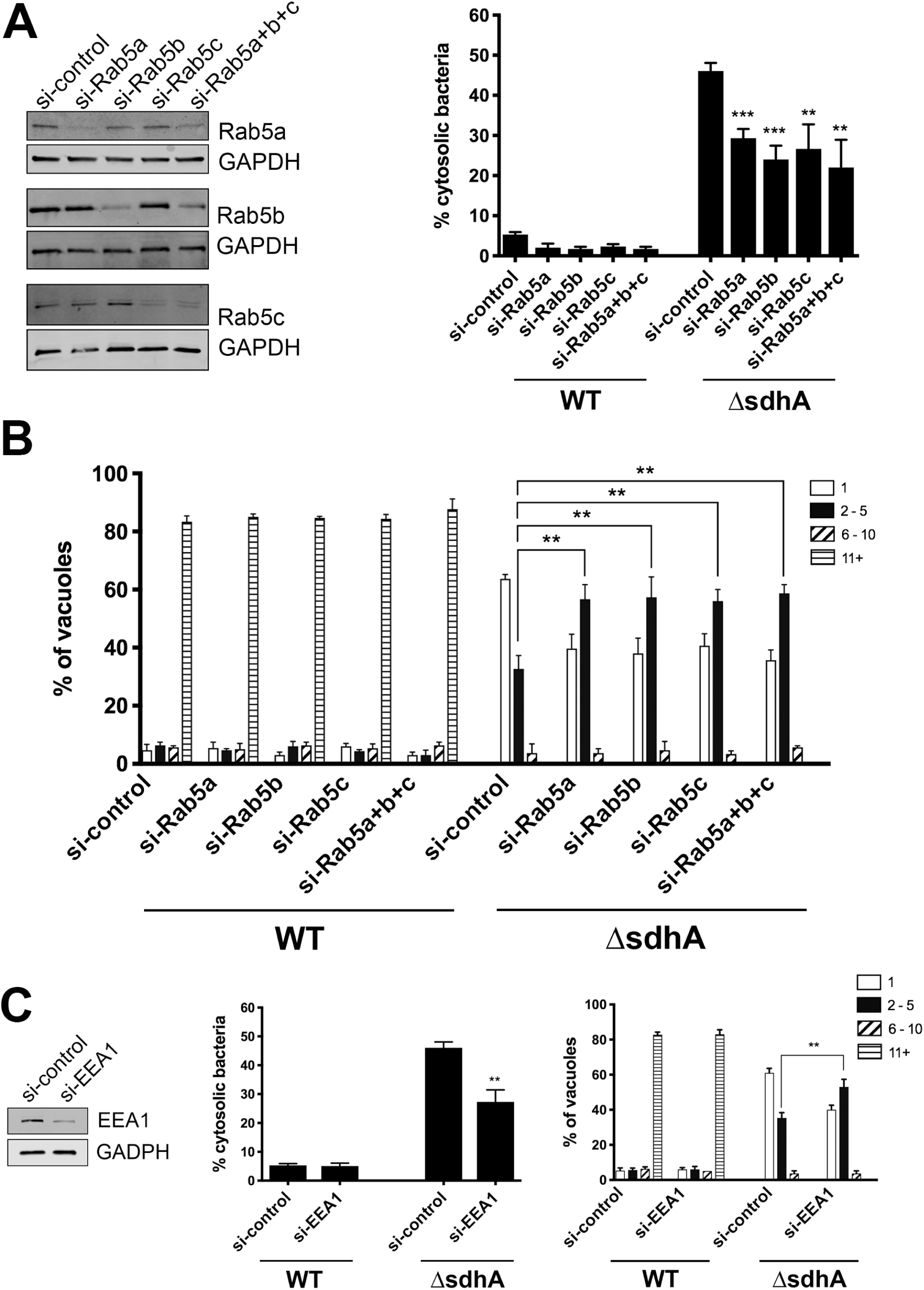
Depletion of proteins that control early endosome dynamics results in increased vacuole integrity after BMDM challenge with *L. pneumophila* Δ*sdhA*. (A) Depletion of Rab5 isoforms partially rescues Δ*sdhA* vacuole integrity. A/J bone marrow-derived macrophages were nucleofected with noted siRNAs. Knockdown efficiency was assessed by immunoblots with noted antibodies (left panels). Nucleofected macrophages were challenged with either WT or Δ*sdhA Legionella*, fixed at 6hpi, and immunostained to determine cytosol exposure. Percent of cytosol-detected bacteria was quantified, as described (Experimental Procedures). **(B)** Growth of noted *L pneumophila* strains in nucleofected BMDM. After BMDM were challenged for 14 h., the cells were fixed and probed to determine yield of bacteria in each macrophage. Number of bacteria per vacuole were quantified microscopically, and binned into 4 groups, displaying the binned vacuole size based on the graph legend (Experimental Procedures; (Luo & Isberg, 2004)). **(C)**. Depletion of downstream Rab5 effector EEA1 rescues Δ*sdhA* vacuole integrity and growth defects. Nucleofection, vacuole integrity and bacterial yields determined as in panels (**A,B**). Statistical analyses were performed on normalized data by unpaired t test (*< 0.05; **<0.01; ***< 0.001; Experimental Procedures).

We next tested the effects on BMDMs that were treated with siRNA against EEA1, a Rab5 effector involved in promoting early endosome vesicle tethering, to determine if depletion of a well characterized target of Rab5 could have similar effects (Christoforidis *et al*., 1999). Depletion of EEA1 in BMDM with an siRNA pool showed lowered steady state levels of the protein that were accompanied by a decrease in the number of cytosol-exposed *L. pneumophila* Δ*sdhA* after 6 hr. infection (**Figure 3C**). Depletion of EEA1 had effects on intracellular growth of the Δ*sdhA* strain that were similar to those observed with Rab5 isoform depletions (**Figure 3B**). These results are consistent with early endosome traffic playing a role in destabilizing the LCV harboring the Δ*sdhA* strain, perhaps because early endosomes are trafficked proximally to the Δ*sdhA* vacuole, resulting in vacuole disruption in the absence of SdhA function.

A model for the action of the SdhA protein is that it acts to limit access of material from early endosomes to the LCV. As EEA1 depletion reduced vacuole permeabilization, and this protein is an important component of early endosomes, the localization of EEA1-positive compartments about LCVs was determined after challenge of macrophages (**Figure 4**). After 4 hours of challenge, BMDMs were fixed and simultaneously stained for EEA1, cytosol-accessible bacteria, total bacteria, and SidC, a marker of the LCV membrane (Luo & Isberg, 2004; Weber *et al*., 2006). EEA1-positive puncta were dispersed throughout the cell, and localization of the puncta about the LCV was determined by image analysis, quantifying the number of colocalization events between EEA1 and the vacuole (**Figure 4A**; Experimental Procedures). The number of EEA1 colocalization events increased significantly at the Δ*sdhA* vacuole compared to the WT vacuole (**Fig. 4A,B**), consistent with EEA1 function playing a role in vacuole destabilization (**Fig. 3C**). Furthermore, this phenotype was rescued by introduction of a plasmid encoding SdhA in the *L. pneumophila* Δ*sdhA* mutant (**Figure 4B,C**). There was no significant correlation between intact or disrupted Δ*sdhA* vacuoles and the EEA1 colocalization, which may be a reflection of transient interactions that cannot be captured by fixed cell assays (**Figure 4D**).

**Figure 4.**
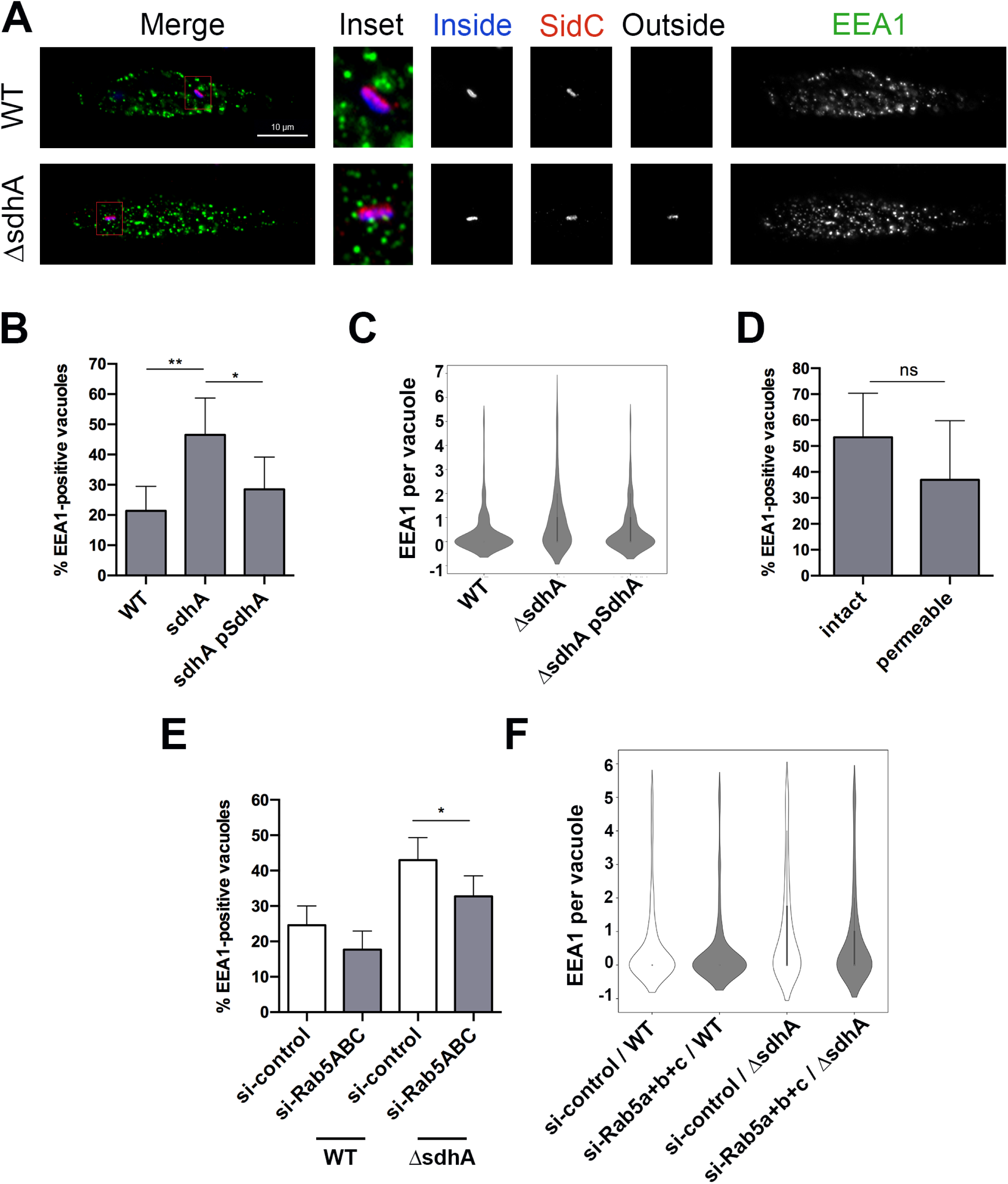
The SdhA protein interferes with contact of early endosomal compartments with the LCV. (A) Representative immunofluorescence microscopy images of the *Legionella*-containing vacuole in infected BMDMs at 4hpi, challenged with noted bacterial strains. Cells were probed with anti-*L. pneumophila* before and after permeabilization, to determine vacuole integrity (Experimental Procedures) as well as anti-SidC and anti-EEA1 after permeabilization. EEA1 and SidC are pseudo-colored in green and red, respectively. Insets are magnified 3.25-fold from the original image by changing resolution. Other panels are identical resolution to original grabbed images. **(B)** Images from (A) were analyzed to quantify colocalization events between the vacuole and EEA1 (Experimental Procedures). Total of 146 vacuoles from 2 experiments were analyzed. **(C)** Frequency distribution of colocalization events from (B). Interquartile range was calculated from integer values. **(D)** Lack of correlation between intact or cytosol-detected (permeable) Δ*sdhA* vacuoles and colocalization of EEA1 at the vacuole. **(E)** BMDMs nucleofected with siRNA were challenged with noted *L. pneumophila* strains and stained as in (A). Images were captured and analyzed after capture to quantify colocalization events of EEA1 at the vacuole) (Experimental Procedures). Total of 182 vacuoles from 2 experiments were analyzed. **(F)** Frequency distribution of colocalization events from (E). Interquartile range was calculated from integer values. Statistical tests were as in Fig. 3.

If aberrant increased localization of EEA1 with the Δ*sdhA* LCV is due to the action of Rab5, then knockdown of Rab isoforms should reverse this effect. As expected, simultaneous knockdown of all three Rab5 isoforms in BMDMS reduced the number of EEA1-positive compartments associated with Δ*sdhA* LCV during infection (**Figure 4E,F**). Therefore, Rab5 isoforms play a role in loss of LCV integrity, causing increased association of EEA1 with the dysfunctional *Legionella* Δ*sdhA*-containing compartment. This result is consistent with proposition that the compartments decorated by EEA1 contribute to this loss of integrity and a function of SdhA protein is to interfere with this traffic.

### A role for membrane egress pathways in destabilizing the Δ*sdhA* mutant

In contrast to Rab5, regulators of anterograde or recycling traffic such as Rab8 and Rab11 have not been associated with restriction of *L. pneumophila* intracellular growth. On the other hand, Rab11a has been connected to regulated disintegration of the *Shigella* vacuole (Mellouk *et al*., 2014; Weiner *et al*., 2016). The identification of knockdown candidates that were downstream effectors of Rab11 (Rab11Fip1; **Figure 2B**) and Rab8 (Vamp3; **Figure 6**) support a role for these proteins in destabilizing the LCV surrounding the Δ*sdhA* mutant (Banerjee *et al*., 2017; Finetti *et al*., 2015; Wilcke *et al*., 2000; Christoforidis *et al*., 1999; Lindsay & McCaffrey, 2004). Surprisingly, only depletion of the Rab11b isoform of Rab11 decreased bacterial exposure to the cytosol after challenge of BMDM with the Δ*sdhA* mutant (**Figure 5A**). Depletion of Rab11a did not rescue the Δ*sdhA* phenotypic defects, while double knockdowns of Rab11a and Rab11b mimicked the defects exhibited by the single depletions that showed rescue (**Figure 5A**). Similarly, only siRNA treatment directed against Rab11b resulted in an increase in the bacterial yield of the Δ*sdhA* mutant within LCVs, arguing that the effects are highly isoform-specific (**Figure 5B**). Similar to the results with Rab5 and other strategies that increase the integrity of the Δ*sdhA* mutant-containing LCV, the stimulation of growth was relatively limited (Creasey & Isberg, 2012).

**Figure 5.**
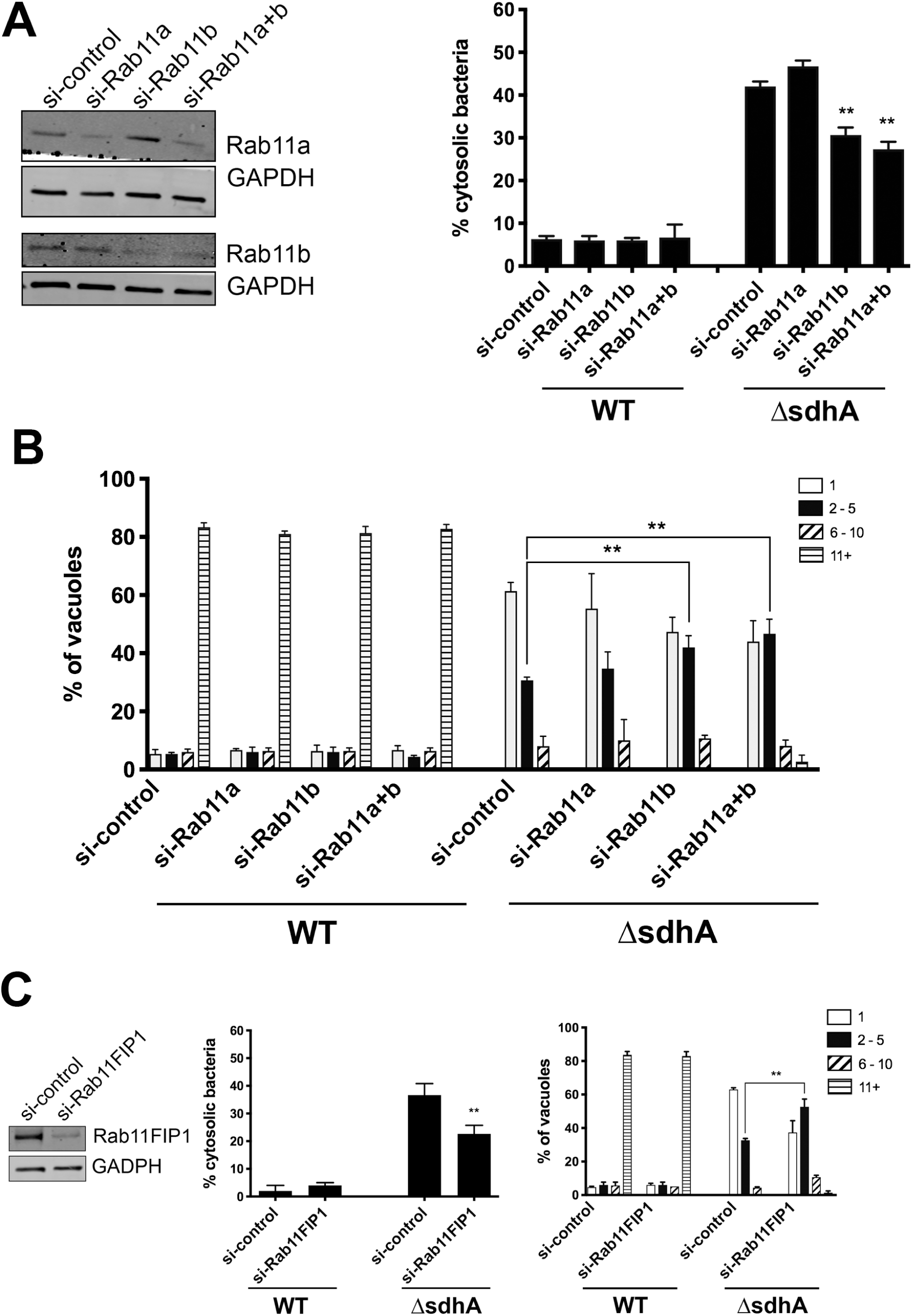
Depletion of proteins associated with recycling dynamics results in increased vacuole integrity after BMDM challenge with *L. pneumophila* Δ*sdhA*. (A) Specific depletion of Rab11b rescues Δ*sdhA* vacuole integrity. A/J bone marrow-derived macrophages were siRNAs treated as in Fig. 3, and knockdown efficiency was assessed (left panels). Nucleofected macrophages were challenged with either WT or Δ*sdhA Legionella* and percent of cytosol-detected bacteria was quantified (Experimental Procedures). **(B)** Growth of noted *L pneumophila* strains in nucleofected BMDM for 14 h. and quantified as described (Fig. 3; Experimental Procedures; (Luo & Isberg, 2004)). **(C)**. Depletion of Rab11 effector Fab11FIP1 rescues Δ*sdhA* vacuole integrity and growth defects. Nucleofection, vacuole integrity and bacterial yields determined as in panels (**A,B**). Statistical analyses were performed on normalized data by unpaired t test (*< 0.05; **<0.01; ***< 0.001; Experimental Procedures).

**Figure 6.**
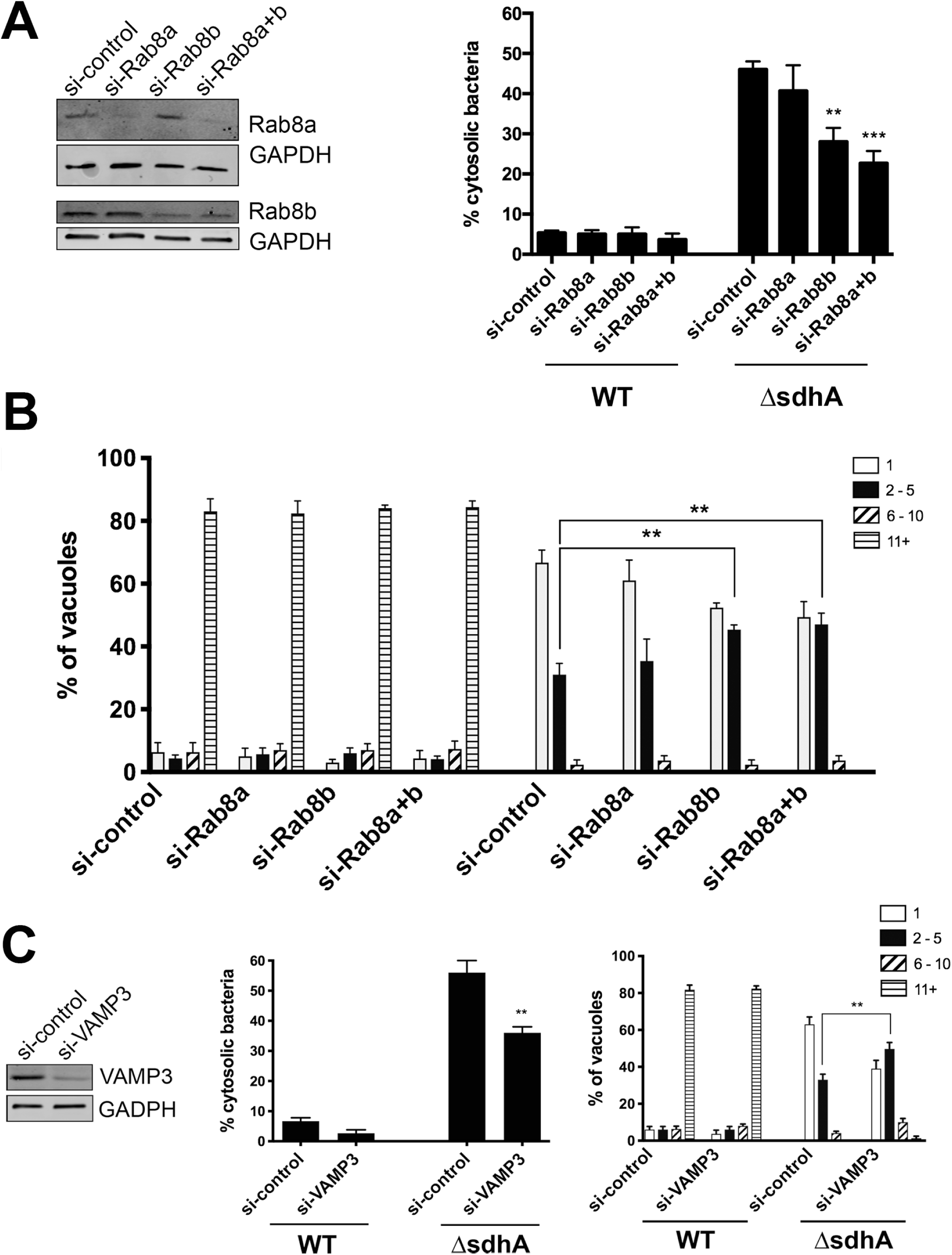
Depletion of anterograde transport Rab8b and a downstream effector reduces bacterial cytosolic exposure. (A) Depletion of Rab8b rescues Δ*sdhA* vacuole integrity. A/J bone marrow-derived macrophages were siRNAs treated and knockdown efficiency was assessed (left panels). Nucleofected macrophages were challenged with either WT or Δ*sdhA Legionella* and percent of cytosol-detected bacteria was quantified (Experimental Procedures). **(B)** Growth of noted *L pneumophila* strains in nucleofected BMDM for 14 h. and quantified as described (Fig. 3; Experimental Procedures; (Luo & Isberg, 2004)). **(C)**. Depletion of Rab8 and Rab11 effector VAMP3 rescues Δ*sdhA* vacuole integrity and growth defects. Nucleofection, vacuole integrity and bacterial yields determined as in panels (**A,B**). Statistical analyses were performed on normalized data by unpaired t test (*< 0.05; **<0.01; ***< 0.001; Experimental Procedures).

From the secondary shRNA screen in BMDMs, we found that silencing Rab11 downstream effector Rab11FIP1 restored Δ*sdhA* vacuole integrity (**Figure 2C**). Nucleofection of BMDMs with siRNA targeting a different region of Rab11FIP1 provided the identical result and demonstrated that depletion of this protein rescued Δ*sdhA* vacuole integrity and growth (**Figure 5C**). Given that several downstream effectors of Rab11 that were targeted by our secondary siRNA screen showed no clear enhancement of LCV integrity (**Figure 2C**), this argues that a specific membrane trafficking pathway promoted by the Rab11B isoform and acting through Rab11FIP1 destabilizes the LCV surrounding the Δ*sdhA* mutant.

Paralleling our results with Rab11, only depletion of the Rab8b isoform showed significant reduction in permeability of the Δ*sdhA* LCV (**Figure 6A**), resulting in increased intracellular growth (**Figure 6B**). When measuring LCV permeability, however, dual depletion with Rab8a and Rab8b increased the statistical significance of this result (**Figure 6A**), indicating that the two proteins may have partial overlap in function. Of particular note regarding the vacuole integrity readouts, both shRNA (Fig. 2B) and siRNA in primary macrophages (Figs 5, 6), gave identical results for both the Rab8 and Rab11 isoforms, further supporting isoform specificity. Similar to knockdown of Rab5 and Rab8 effectors, siRNA directed against VAMP3, a SNARE protein that promotes vesicle fusion events and is a downstream effector of both Rab11 and Rab8, was sufficient to partially restore Δ*sdhA* vacuole integrity and intracellular growth (Banerjee *et al*., 2017; Finetti *et al*., 2015; Wilcke *et al*., 2000; Christoforidis *et al*., 1999; Lindsay & McCaffrey, 2004) (**Figure 6C**).

Based on these findings, we expect that membrane material driven by these GTPases will be closely associated with the Δ*sdhA* LCV, contributing to vacuole disruption. In the presence of SdhA function, in contrast, access to these compartments should be limited. To this end, the localization of the Rab11 effector FIP1 was analyzed, using the same assay employed to detect EEA1 localization (**Figure 4**). BMDM challenged for 4 hrs with either the WT or Δ*sdhA* strain were fixed and probed for vacuole permeability and association of Rab11FIP1 with the LCV was determined (**Figure 7**). Antibody against Rab11FIP1 labeled distinct puncta dispersed throughout the cell (**Figure 7A**). Similar to EEA1, Rab11FIP1 colocalized at a higher frequency with the Δ*sdhA* vacuole compared to the WT vacuole, with the phenotype rescued by complementation of SdhA on plasmid *in trans* (**Figure 7B,C**). Again, we observed no significant correlation between intact or disrupted Δ*sdhA* vacuoles and Rab11FIP1 colocalization (**Figure 7D**). Similar to the results obtained with Rab5 and EEA1 localization, knockdown of the Rab11b isoform reduced the number of Rab11FIP1 compartments associated with the Δ*sdhA* vacuole (**Figure 7E,F**). This argues that in the absence of SdhA function, the heightened level of Rab11FIP1-staining material associated with the LCV is a consequence of Rab11b trafficking events that result in vacuole disruption.

**Figure 7.**
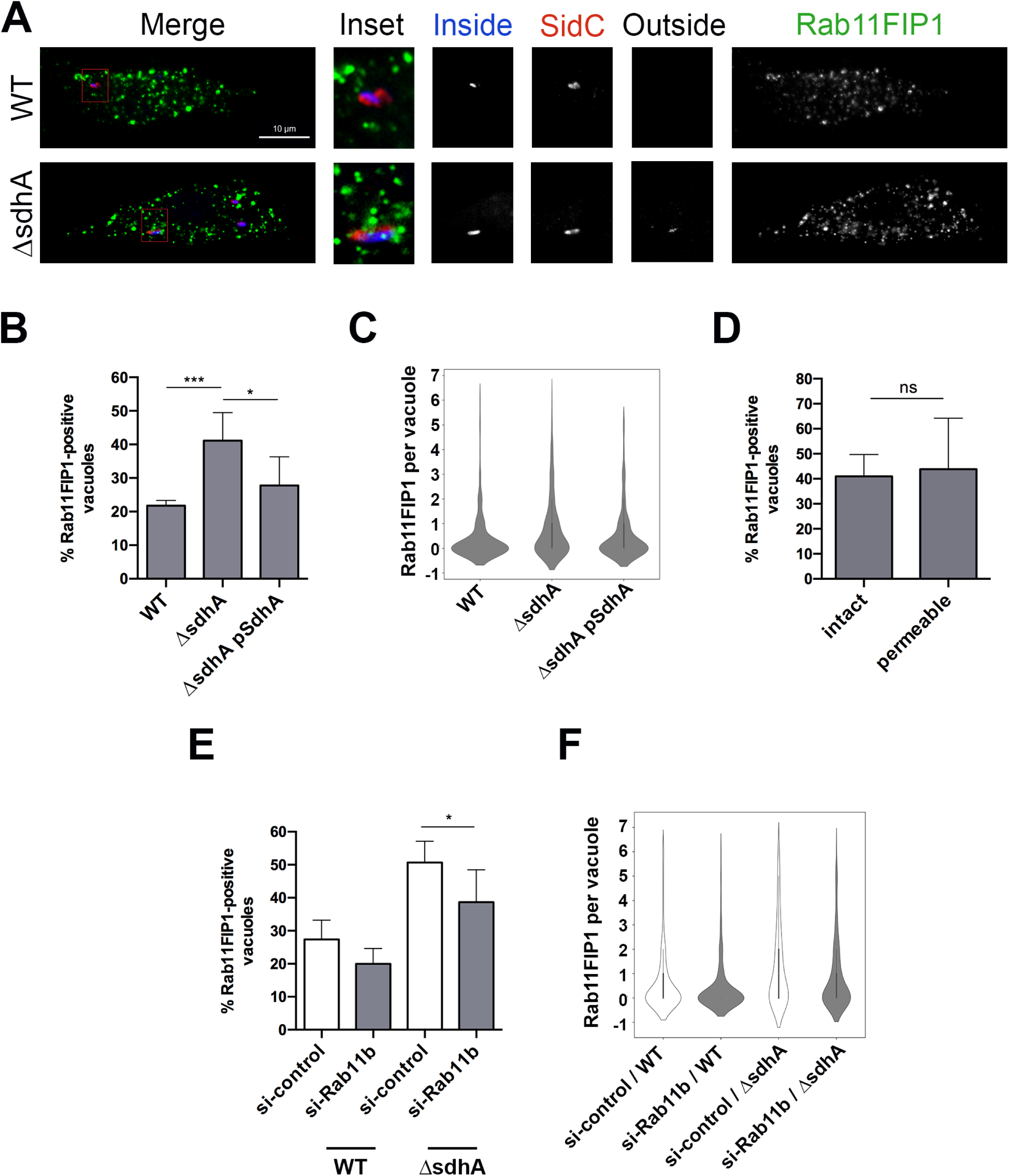
SdhA functions to interfere with Rab11FIP1-LCV interactions. (A) Representative immunofluorescence microscopy images of the *Legionella*-containing vacuole in infected BMDMs at 4hpi, challenged with noted bacterial strains. Cells were probed with anti-*L. pneumophila* before and after permeabilization, to determine vacuole integrity (Experimental Procedures) as well as anti-Rab11FIP1 and SidC after permeabilization. Rab11FIP1 and SidC are pseudo-colored in green and red, respectively. Insets are magnified 3.25-fold from the original image by changing resolution. Other panels are identical resolution to original grabbed images. **(B)** Images from (A) were analyzed in Volocity to quantify colocalization events between the vacuole and Rab11FIP1. Total of 181 vacuoles from 2 experiments were analyzed. **(C)** Frequency distribution of colocalization events from (B). Interquartile range was calculated from integer values. **(D)** Lack of correlation between intact or cytosol-detected (permeable) Δ*sdhA* vacuoles and colocalization of Rab11FIP1 at the vacuole. **(E)** BMDMs were nucleofected with siRNA, were challenged with noted *L. pneumophila* strains and stained as in (A). Images were captured and analyzed after capture to to quantify colocalization events of Rab11FIP1 at the vacuole (Experimental Procedures). Total of 158 vacuoles from 2 experiments were analyzed. **(F)** Frequency distribution of colocalization events from (E). Interquartile range was calculated from integer values.

## Discussion

In this study, screens were performed that identified host membrane trafficking factors responsible for causing vacuole disruption and interfering with intracellular growth of an *L. pneumophila* Δ*sdhA* strain. We initially used a luciferase reporter assay to positively select for siRNA-depleted cells that showed increased intracellular growth of the mutant strain. Overall, 20 candidate genes were identified after screening 112 genes. To demonstrate specificity for knockdowns that result in increased LCV integrity relative to that observed in bone marrow-derived macrophages (BMDMs), we turned to high-throughput microscopy analysis to identify factors specifically involved in antagonizing vacuole integrity of the *L. pneumophila* Δ*sdhA* mutant. From a primary shRNA screen introduced into BMDMs by lentivirus, we identified 8 candidate genes, with the majority being Rab GTPases involved in endocytic and membrane-recycling. Genes of particular interest were Rab5, Rab8, and Rab11. Strikingly, Rab11a had been previously shown to regulate vacuole rupture as an early event that supports intracellular growth of *Shigella* after uptake into cultured cells (Mellouk *et al*., 2014; Weiner *et al*., 2016). This raises the possibility that trafficking events that support intracellular growth of a cytosolic pathogen (*S.flexneri*) interfere with the biogenesis of the membrane surrounding an intravacuolar organism (*L. pneumophila*).

Validation of screen hits revealed that rescue of Δ*sdhA* vacuole integrity and growth was specific to depletion of individual isoforms. One exception to this observation was silencing Rab5a, Rab5b, or Rab5c, in which we found that individual depletion of each isoform was sufficient to reduce vacuole disruption (**Figure 3A**). This phenotype is consistent with previous studies demonstrating that the three isoforms localize to the same endocytic compartment and cooperatively regulate endocytic events (Bucci *et al*., 1995). Depletion of all three isoforms rescued the Δ*sdhA* phenotype to a similar degree as the depletion of individual isoforms, indicating that the isoforms function collectively rather than separately during trafficking.

In contrast to the situation with the Rab5 isoforms, specific silencing of Rab11b, but not Rab11a, enhanced the integrity of the Δ*sdhA* vacuole (**Figure 5A**). Furthermore, knockdown of both Rab11 isoforms produced a similar phenotype as individual knockdown of Rab11b, confirming that only Rab11b is required for Δ*sdhA* vacuole disruption. It has been shown in other cell types that these two isoforms can localize to distinct compartments within the cell, which suggests a division of function between the two isoforms that is linked to their association with different compartments (Kelly *et al*., 2012; Lapierre *et al*., 2003). Of particular note in this regard is that the Rab11a isoform specifically supports growth of *S. flextneri*. The isoform differences between the two pathogens may reflect spatial and temporal differences in the biogenesis and degradation of the bacterium-containing compartments. After uptake into host cells, depletion of Rab11a interferes with degradation of the *Shigella-*containing vacuole, while Rab11b has no documented effect. As degradation of the *Shigella-*containing vacuole occurs shortly after bacterial uptake at the periphery of cells, Rab11a may drive compartment destabilization in this region. In contrast, *Legionella* vacuole degradation occurs hours after infection, allowing access to perinuclear regions that could allow an interface with Rab11b-containing compartments.

A similar phenomenon was discovered when individual Rab8 isoforms were depleted in primary macrophages. Silencing of Rab8b, but not Rab8a, increased the stability of vacuoles harboring the Δ*sdhA* strain and resulted in enhanced growth for the mutant (**Figure 6A**). Very little is known about the difference between the two Rab8 isoforms, although it is postulated that they function in both distinct and as well as overlapping compartments in various cell types (Chen *et al*., 2001; Sato *et al*., 2014). Unlike the situation with Rab11a/b, experimental knockdown of Rab8a in combination with Rab8b enhances rescue relative to individual depletion of the Rab8b isoform. This additive effect is consistent with a small pool of Rab8a that has either overlapping function or overlapping localization with Rab8b, resulting in a fraction of the Δ*sdhA*-containing vacuoles being destabilized by the Rab8a isoform.

We investigated if downstream effectors of the Rab GTPases were involved in facilitating Δ*sdhA* vacuole disruption. We demonstrated that depletion of the Rab5 effector EEA1 and the Rab11b effector Rab11FIP1 partially restored integrity of the Δ*sdhA-*containing vacuole and increased intracellular growth of the mutant, generating phenotypes indistinguishable from depletion of the GTPases (**Figure 3C**,5C). We also found that depletion of VAMP3, a downstream regulator fusion events controlled by both Rab8b and Rab11b, could result in an increased number of intact vacuoles harboring the Δ*sdhA* strain (**Figure 6C**) (Banerjee *et al*., 2017; Finetti *et al*., 2015; Wilcke *et al*., 2000). VAMP3 is SNARE protein that drives membrane fusion, thus pointing to a model in which disruption of membrane integrity is a consequence of fusion with a compartment that destabilizes the LCV (Hu *et al*., 2007).

The effectors implicated in vacuole destabilization are consistent with their defining either two different types of compartments that can destabilize the LCV or identifying distinct steps in the destabilization process. We found that depletion of Rab5 isoforms or Rab11b decreased the percentage of EEA1 and Rab11FIP1 localization at the Δ*sdhA* vacuole, respectively. EEA1 is involved in *tethering* Rab5-positive endosomes to membranes enriched in PI(3)P while Rab11FIP1 is known to be involved in *docking* recycling vesicles to membranes enriched in PI(3,4,5,)P_3_ (Christoforidis *et al*., 1999; Lindsay & McCaffrey, 2004). Therefore, the nature of the lipid components could define different compartments, or these proteins could define tethering (EEA1), docking (Rab11Fip1), and fusion (Vamp3) events. Consistent with either model, we found that EEA1 and Rab11FIP1 localized to the Δ*sdhA*-containing vacuole at a higher frequency compared to the WT-containing vacuole, indicating that they may play a direct role in destabilizing the vacuole harboring the mutant (**Figure 4**, 7). One of the striking properties of the LCV is that is appears to be walled off from the host early endosomal trafficking system (Isberg *et al*., 2009). The presence of EEA1 associated with events leading to vacuole rupture indicates that constructing of such a firewall is essential for preserving vacuole integrity.

The mechanical details of Δ*sdhA* vacuole disruption remain unclear. Surprisingly, neither EEA1 nor Rab11FIP1 recruitment showed any preference for intact or disrupted vacuoles harboring the Δ*sdhA* strain. Vacuole interactions with destabilizing compartments could be transient, making it difficult to distinguish differences based on single time point assays. Alternatively, if a variety of membrane compartments can destabilize the LCV, probing for these two proteins will not capture the total spectrum of disruptive compartments. Of note is the possibility that the increased frequency of EEA1 and Rab11FIP1 at the Δ*sdhA* vacuole compared to WT identifies a phospholipid motif that signals for self-destruction of the compartment by a number of membrane trafficking events. Consistent with the formation of a unique lipid environment in response to the *sdhA* mutant, the compartment is particularly sensitive to the action of the *L. pneumophila* lysophospholipase PlaA. (Creasey & Isberg, 2012). Presumably, lipids are found on this vacuole that are missing from the WT, in turn serving as substrates for PlaA.

Not considered here is the possibility that cargo being delivered to the Δ*sdhA* vacuole from the endocytic-recycling pathway directly destroys membranes. Delivery of toxic cargo to the bacteria-containing vacuole has been observed in other pathogens, such as *Salmonella* (McGourty *et al*., 2012). It has been demonstrated in macrophages that Rab11 traffics the NADPH oxidase flavocytochrome B, and it is possible that this cargo could be delivered to the Δ*sdhA* vacuole resulting in oxidative destabilization (Casbon *et al*., 2009). It is interesting to note that vacuole degradation is an essential step necessary to release *S. flexneri* into the cytosol and initiate intracellular replication. These events appear to require recruitment of Rab11A-containing vesicles that could carry destabilizing cargo (Mellouk *et al*., 2014; Weiner *et al*., 2016). Therefore, an event that for one pathogen is toxic, for another is an essential step in the biogenesis process leading to intracellular growth.

Altogether, this study describes a genetic screen that identified a specific set of host factors that are required for Δ*sdhA* vacuole disruption. Future studies should elucidate the exact mechanism by which these proteins facilitate vacuole disruption and the nature of the cargo being delivered to the vacuole that destabilizes the vacuole.

## Experimental Procedures

### Bacterial culture and media

All *Legionella pneumophila* strains used in this study are derived from the Philadelphia 1 isolate and are described in Table 2 (Berger & Isberg, 1993). Luminescent *L. pneumophila* was constructed using *P_ahpC_*::*luxCDABE* as previously described (Coers *et al*., 2007; Ensminger *et al*., 2012). *L. pneumophila* strains were grown on plates containing charcoal and yeast extract buffered with ACES [N-(2-acetamido)-2-aminoethanesulfonic; Sigma] adjusted to pH 6.9 and supplemented with 0.4 mg/mL of _L_-cysteine and 0.135 mg/mL of ferric nitrate (CYE), as well as 0.1 mg/mL thymidine and/or kanamycin (Sigma) when necessary. Liquid cultures of *L. pneumophila* were prepared in the same medium, but without charcoal and agar (AYE) (de Jesus-Diaz *et al*., 2017). Overnight *L. pneumophila* cultures were prepared by inoculating AYE broth with a bacterial patch and serially diluting cultures 1 to 2. Liquid cultures were incubated overnight at 37°C with shaking. For infections, overnight cultures were used, and all strains were grown to post-exponential phase (*A_600_* of 3.5 to 4.0) (Byrne & Swanson, 1998). The approximate multiplicities of infection were determined by assuming that an *A_600_* = 1.0 is equivalent to 10^9^ bacteria/mL.

**Table 2:** Strains used in this study

| Strain/Plasmid | Genotype | Description | Reference |
| --- | --- | --- | --- |
| WT | Lp02, Philadelphia-1, rpsL<br>hsdR thyA- | wild-type strain | [45] |
| ΔsdhA | Lp02 ΔsdhA | SdhA deletion | [12] |
| ΔsdhA lux+ | Lp02 ΔsdhA kan<sup>R</sup> P<sub>ahpC</sub>::lux | SdhA deletion, Lux+ | This study |
| pSdhA | pJB908*sdhA* | Expression vector for<br>SdhA | [12] |

### Mammalian cell culture

RAW 264.7 cells (ATCC TIB-71) were grown in Dulbecco modified Eagle medium (DMEM, Gibco) supplemented with heat-inactivated fetal bovine serum (FBS). Bone marrow-derived macrophages (BMDMs) were isolated from femurs and tibias of female A/J mice as previously described and frozen in heat-inactivated FBS with 10% dimethyl sulfoxide (DMSO) (Auerbuch *et al*., 2009). BMDMs were plated in RPMI 1640 (Gibco) supplemented with 10% heat-inactivated FBS (Invitrogen) and 1mM _L_-glutamine (Gibco) one day prior to challenge with *L. pneumophila*. Animal protocols were approved by the Institutional Animal Care and Use Committee of Tufts University.

### Primary siRNA library screen

The Mouse siGENOME siRNA Library of 112 siRNA directed against membrane trafficking genes (Dharmacon, Lafayette, CO; GU-015500) was tested in a high-throughput screen. The library consists of a pool of four different oligos for each target gene. RAW 264.7 cells were seeded overnight in RPMI medium (Thermo Fisher Scientific) containing 10% (vol/vol) FBS at a density of 1.25 x 10^4^ cells per well in a 96 well white-bottom plate (Thermo Fisher Scientific). The next day, the cells were transfected with 50nM siRNA using Lipofectamine RNAiMAX Reagent (Thermo Fisher Scientific). The screen was designed with three negative control wells, transfected with non-targeting siRNA, and one well for each experimental siRNA target. After 24 hours of transfection, the medium was replaced to reduce cell cytotoxicity. After 48 hours of transfection, cells were challenged at an MOI = 0.5 with *L. pneumophila* Δ*sdhA* Lux ^+^ grown to post-exponential phase. At 1 hr post-challenge, extracellular bacteria were removed by washing in PBS and cells were incubated at 37°C, 5% CO_2_ in Phenol Red-free RPMI media containing 10% FBS. Relative light units (RLU) of each well were measured every 12 hours in a Tecan M200 Pro plate reader. A total of 8 plates were screened to collect at least three replicate measurements for each gene target. RLU from each experimental well was normalized to the average RLU of the negative control wells on the same plate. Using normalized data, Z_MAD_ scores (equivalent the number of median absolute deviations (MAD) from the median) were calculated for each experimental siRNA target and were used for selection of hits (Chung *et al*., 2008). The classification of the scale of the siRNA effects based on the ZMAD score was defined as follow: | ZMAD | ≥ 2 for extremely strong RLU, 2 > | ZMAD | ≥ 1.5 for very strong RLU, 1.5 > | ZMAD | ≥ 1 for strong RLU, 1 > | ZMAD | ≥ 0.5 for moderately strong RLU, 0.5 > | ZMAD | ≥ 0 for no effect, 0 > | ZMAD | ≥ -0.5 for moderately weak RLU, -0.5 > | ZMAD | ≥ −1 for weak RLU, −1 > | ZMAD | ≥ −1.5 for very weak RLU, −1.5 > | ZMAD | for extremely weak RLU. Only siRNAs that had a ZMAD of ≥ 1.5 were considered of interest for further analysis in the secondary shRNA library screen, as described next.

### Secondary shRNA library screen

Selected hits from the primary siRNA library screen and additional related targets were tested in a screen using shRNA constructions introduced into BMDMs. The shRNA were obtained from the Broad Institute Genetic Pertubation Platform (https://www.broadinstitute.org/genetic-perturbation-platform). To this end, lentiviral vector library plates were constructed with three negative control shRNAs against LacZ and 92 experimental lentiviral shRNAs against 23 target genes (four shRNAs per target). All shRNAs were contained on third-generation transfer plasmid pLKO.1, which confers puromycin resistance. On day 0, progenitor cells derived from the bone marrow of A/J mice (Experimental Procedures) were seeded at a density of 5 x 10^4^ cells per well in a 96 well clear-bottom black plate (Corning) in RPMI medium containing 30% (vol/vol) L-cell supernatant, 20% FBS, and 1% penicillin-streptomycin (BMM media, Gibco). On day 4, polybrene (Sigma) and 2×10^5^ infectious units of lentivirus were added to the wells, and transduction was initiated by centrifugation at 2200 rpm at 37°C for 20 minutes. On day 5, conditioned medium was replaced to reduce cell cytotoxicity. On day 6, transductants were selected in conditioned medium containing 3µg/mL puromycin. On day 8, the supernatants were removed and replaced with puromycin-free conditioned medium to recover transduced clones. On day 10, the cultured medium was changed to RPMI containing 10% FBS. Terminally differentiated macrophage transductants were challenged with *L. pneumophila* Δ*sdhA* at MOI = 1 for 1 hr, followed by removal of extracellular bacteria by washing in PBS. After 6 hours, infected macrophages were fixed with 4% (vol/vol) paraformaldehyde in PBS and disrupted vacuoles were stained as described in “Vacuole Integrity & Intracellular Replication Assay.” The screen was performed on triplicate shRNA library and plates were stored in PBS at 4°C until analysis.

After fixation and staining, images of each well were captured using ImageXpress (Molecular Devices) (Huang *et al*., 2011). All images were captured using 20X Plan Apo lens. Images were captured with the FITC (ex. 490, em. 525) and Texas Red (ex. 590, em. 617) filter sets. Sixteen images were captured at the center of each well. A total of three plates were screened to acquire three replicates for each knockdown condition. The “Cell Scoring” function from the MetaXpress software was used to calculate the number of *Legionella*-containing vacuoles (LCV) in each channel per image. LCVs were defined as having a pixel intensity of at least 50 gray levels above background, with a minimum width of 6 pixels and a maximum width of 12 pixels. LCVs were scored as disrupted if the pixel intensity in the FITC channel overlapped with the pixel intensity in the Texas Red channel. The number of disrupted LCVs in each well were recorded and analyzed in Microsoft Excel. Raw data from each experimental well was normalized to the average of the negative control wells. Statistical analyses were performed on normalized data by unpaired t test (*P < 0.05).

### Nucleofection

Frozen differentiated BMDMs were recovered and plated at a density of 5×10^6^ cells in a 10cm dish (Falcon) in RPMI medium containing 10% (vol/vol) FBS and 10% (vol/vol) L-cell conditioned supernatant. The next day, cells were lifted in cold PBS (Gibco) and resuspended in RPMI medium containing FBS (Swanson & Isberg, 1995). Resuspended cells were aliquoted into 1.5 ml microfuge tubes at 1×10^6^ cells per tube and pelleted by centrifugations for 10 minutes at 200xg. Cell pellets were resuspended in nucleofector buffer (Amaxa Mouse Macrophage Nucleofector Kit (Lonza)) and 2ug of siRNA was added to each tube (Dharmacon). Cells were transferred to a cuvette and nucleofected in the Nucleofector 2b Device (Lonza) under Y-001 program settings. Nucleofected macrophages were immediately recovered in RPMI medium containing 10% FBS and 10% L-cell supernatant. Cells were plated in 8-well chamber slides (Millipore Sigma) for microscopy assays or in 12-well plates (Corning) to prepare extracts for immunoblotting.

### Immunoblotting

Efficiency of siRNA silencing in nucleofected cells was determined by immunoblot. Nucleofected macrophages were plated in 12-well plates in RPMI medium containing 10% (vol/vol) FBS and 10% (vol/vol) L-cell supernatant. After 48 hours, cells were lysed using 4X SDS Laemmli sample buffer (Bio-Rad) and boiled for 5 minutes. After fractionation on SDS-polyacrylamide gels (Bio-Rad), proteins were transferred to nitrocellulose membranes, blocked in 4% (vol/vol) milk in TBST [0.05 M Tris buffered saline (NaCl = 0.138 M, KCl = 0.0027 M); (Tween-20 = 0.05%, pH 8.0) (Sigma-Aldrich) and probed with antibodies to Rab5A (Cell Signaling, 1:500), Rab5B (Santa Cruz Biotechnology, 1:500), Rab5C (Novus Biologicals, 1:500), Rab11A (Cell Signaling, 1:500), Rab11B (Cell Signaling, 1:500), Rab8A (Cell Signaling Technologies, 1:500), Rab8B (Proteintech, 1:500), EEA1 (BD Biosciences, 1:500), Rab11FIP1 (Cell Signaling, 1:500), VAMP3 (Synaptic Systems, 1:500) or GAPDH (Santa Cruz Biotechnology, 1:1000) in 4% milk/TBST. Immunoblotting with primary antibodies was carried overnight at 4°C. After washing in TBST, secondary antibodies Dylight anti-rabbit IgG 680, Dylight anti-mouse IgG 680, Dylight anti-rabbit IgG 800, and Dylight anti-mouse IgG 800 (Cell Signaling 1:20,000) were incubated in 4% milk/TBST for 45 minutes at room temperature. Capture and analysis was performed using the Odyssey Scanner and the Image Studio software (LI-COR Biosciences).

### Vacuole Integrity & Intracellular Replication Assays

For fluorescence microscopy experiments, nucleofected BMDMs were seeded in 8-well chamber slides (Millipore Sigma) at a density of 5 x10^4^ cells per well in RPMI medium containing 10% (vol/vol) FBS and 10% (vol/vol) L-cell supernatant. After 48 hours, the medium was changed to RPMI containing 10% FBS only. Nucleofected macrophages were challenged with post-exponential phase *L. pneumophila* strains at a MOI = 1. At 1h after challenge, extracellular bacteria were removed by washing in PBS, and at the indicated times, macrophages were fixed with 4% (vol/vol) paraformaldehyde in PBS and blocked in PBS containing 4% BSA overnight at 4°C.

For detection of *L. pneumophila* disrupted vacuoles, nucleofected BMDMs were incubated with rabbit anti-*L. pneumophila* antisera (1:20,000) for 1h at 37°C in PBS containing 4% BSA followed by anti-rabbit IgG Alex Fluor 488 (Invitrogen, 1:500) for 1h at 37°C in PBS containing 4% BSA. After washing three times in room temperature PBS, cells were permeabilized with ice-cold methanol for 20 seconds, blocked in PBS containing 4% BSA for 25 minutes at room temperature, and incubated again with *L. pneumophila* antisera (1:20,000) in PBS containing 4% BSA for 1h at 37°C, followed by anti-rabbit IgG Alex Fluor 594 (Invitrogen, 1:500) for 1h at 37°C in PBS containing 4% BSA. Disrupted vacuoles were identified by the simultaneous staining of bacteria with both Alexa Fluor 488 (bacteria detected prior to permeabilization) and Alexa Fluor 594 (bacteria detected after permeabilization) antibodies by fluorescence microscopy. For analysis of *L. pneumophila* intracellular replication at a single-cell level, nucleofected BMDMs were permeabilized with ice-cold methanol for 20 seconds, blocked in 4% BSA for 25 minutes at room temperature, and then incubated with anti-*L. pneumophila* sera (1:20,000) for 1h at 37°C, followed by anti-rabbit IgG Alex Fluor 488 (1:500) for 1h at 37°C. The number of bacteria contained in each vacuole was quantified by fluorescence microscopy (Creasey & Isberg, 2012).

### Localization of EEA1 or Rab11FIP1 at the *Legionella* vacuole

For EEA1 or Rab11FIP1 localization studies, BMDMs were challenged with *L. pneumophila* strains in 8-well chamber slides. After 4 hours of incubation, cells were fixed with 4% paraformaldehyde in PBS and blocked in PBS containing 4% BSA for 1 hour at room temperature. For EEA1 staining, cells were incubated overnight at 4°C with anti-mouse EEA1 antibody (BD Biosciences, 1:50) in PBS containing 4% BSA. The next day, cells were incubated with anti-mouse IgG Alexa Fluor 488 (Invitrogen, 1:500) for 1h at 37°C in PBS containing 4% BSA. Cells were incubated with rabbit anti-*L. pneumophila* (1:20,000) for 1h at 37°C in PBS containing 4% BSA, followed by anti-rabbit IgG Dylight 405 (Invitrogen, 1:200) for 1h at 37°C in PBS containing 4% BSA. Cells were permeabilized with ice-cold methanol for 20 seconds, blocked in PBS containing 4% BSA for 25 minutes at room temperature, and incubated with rat anti-SidC antibody (1:200) for 1h at 37°C in PBS containing 4% BSA, followed by anti-rat IgG Alex Fluor 594 antibody (Invitrogen, 1:200) in PBS containing 4% BSA. Cells were incubated with *L. pneumophila* antisera (1:20,000) for 1h at 37°C in PBS containing 4% BSA, followed by anti-rabbit IgG Alex Fluor 647 (Invitrogen, 1:500) for 1h at 37°C in PBS containing 4% BSA. For Rab11FIP1 staining, cells were incubated overnight at 4°C with anti-rabbit Rab11FIP1 antibody (Cell Signaling Technologies) and staining procedures followed as described above. Chamber slides were mounted with ProLong Gold (Life Technologies) and imaged with a 63X objective lens, using a Zeiss Axio Observer.Z1 (Zeiss) fluorescent microscope, an Apotome.2 (Zeiss), and an ORCA-R^2^ digital CCD camera (Hamamatsu). Images were acquired under Cy5 (ex. 650, em. 665), FITC (ex. 490, em. 525), Texas Red (ex. 590, em. 617), and DAPI (ex. 346, em. 442) filter sets, and 2.5µm z stacks were acquired per image (10 steps were imaged at 0.25µm step size). Three replicates for each strain were imaged.

Volocity image analysis software (Quorum Technologies) was used to measure colocalization between EEA1 or Rab11FIP1 and the *Legionella*-containing vacuole (LCV). Objects were gated based on signal intensity as the following: bacteria were thresholded on Cy5 intensity, SidC was thresholded on Texas Red intensity, EEA1 or Rab11FIP1 was thresholded on FITC intensity. The bacteria and SidC gates were combined to define the LCV gate. The numbers of disrupted vacuoles were manually quantified in the DAPI channel. The number of EEA1 or Rab11FIP1 objects overlapping in the LCV gate were quantified and correlated with the number of disrupted vacuoles for each image. Each experiment contained three replicates for each *Legionella* strain, and quantification of ∼30 vacuoles for each *Legionella* strain. Statistical analyses were performed on each replicate by unpaired t test (*P < 0.05, **P < 0.01, ***P<0.001).

## Supporting information

Supplemental Data complete

## Acknowledgments

This work was supported by NIH/National Institute of Allergy and Infectious Diseases (NIAID) Training Grant T32GM07310 to I.A. as well as by HHMI and NIAID grant R01AI113211 to R.R.I. We thank Dr. Albert Tai from Tufts University Core Facility for technical assistance in screens and Dr David Root of the Broad Institute for consultations regarding shRNA experiments. We thank members of the Isberg lab for review of the manuscript.

## Author contributions

A.I, W.Y. and R.R.I. designed research; A.I. performed research; A.I, W.Y. and R.R.I. wrote the paper.

