## Supplemental Data complete for "Components of the endocytic and recycling trafficking pathways interfere with the integrity of the *Legionella*-containing vacuole"

### Supplemental Figure 1

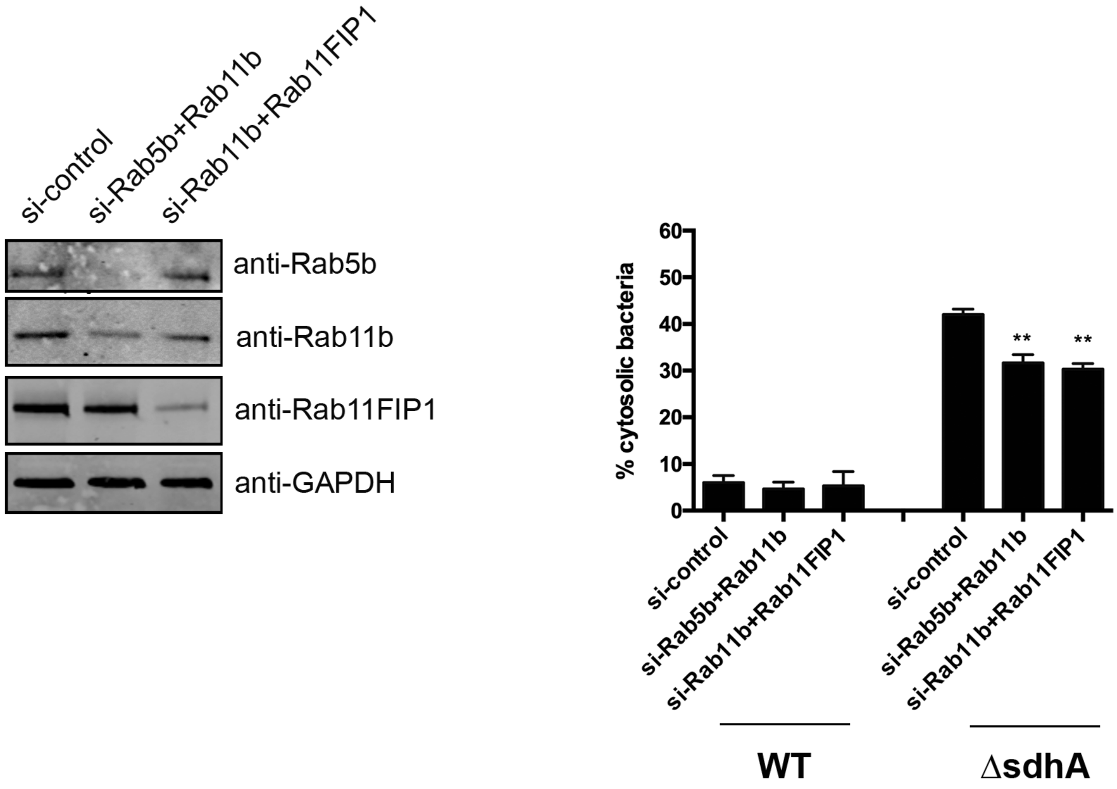

**Supplemental Table 1. Results of high-throughput siRNA screen for enhanced intracellular growth of *L. pneumophila* Δ*sdhA* Lux$^{+}$.**

**ZMAD Value Key**

-2 -1.5 -1 -0.5 0 0.5 1 1.5 2

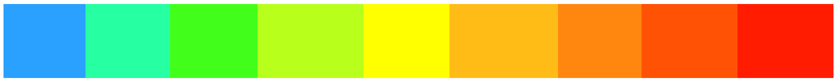

| | | 12 hpi | | 24 hpi | |
|---|---|---|---|---|---|
| **Gene** | **Accession** | **ZMAD value** | **Code** | **ZMAD value** | **Code** |
| DNM3 | NM_172646 | 0.32 | | 0.61 | |
| ACTR2 | NM_146243 | 0.19 | | -0.15 | |
| ACTR3 | NM_023735 | 1.38 | | 1.05 | |
| ADAM10 | NM_007399 | -1.40 | | -2.46 | |
| AI642036 | NM_001045520 | 0.01 | | 0.28 | |
| AMPH | NM_175007 | 0.28 | | -0.70 | |
| AP1B1 | NM_007454 | -0.43 | | -1.03 | |
| AP1M1 | NM_007456 | -1.49 | | -2.23 | |
| AP1M2 | NM_009678 | -0.62 | | -0.54 | |
| AP2A1 | NM_001077264 | -1.20 | | -2.29 | |
| AP2A2 | NM_007459 | -0.78 | | -0.73 | |
| AP2B1 | NM_027915 | 0.67 | | -0.26 | |
| AP2M1 | NM_009679 | -0.17 | | 0.09 | |

| | | 12 hpi | | 24 hpi | |
|---|---|---|---|---|---|
| **Gene** | **Accession** | **ZMAD value** | **Code** | **ZMAD value** | **Code** |
| ARF1 | NM_007476 | -0.81 | | -0.89 | |
| ARF6 | NM_007481 | -0.78 | | -1.18 | |
| ARFIP2 | NM_029802 | 0.68 | | 2.40 | |
| ARPC1B | NM_023142 | -0.99 | | -0.84 | |
| ARPC2 | NM_029711 | -0.12 | | 0.30 | |
| Arpc3 | NM_019824 | 0.62 | | 0.10 | |
| ARPC4 | NM_026552 | 1.34 | | -0.40 | |
| Arrb1 | NM_178220 | 2.72 | | 2.17 | |
| ARRB2 | NM_145429 | 1.66 | | 0.91 | |
| ATM | NM_007499 | -0.73 | | -1.22 | |
| ATP6V0A1 | NM_016920 | -0.18 | | -0.29 | |
| BIN1 | NM_001083334 | 0.69 | | 0.53 | |
| C730014M21RIK | XM_356089 | 1.09 | | -0.03 | |
| CAMK1 | NM_133926 | 1.23 | | 0.54 | |
| CAV | NM_007616 | -0.68 | | -0.14 | |
| CAV2 | NM_016900 | 0.46 | | 0.42 | |
| CAV3 | NM_007617 | -0.07 | | -0.73 | |
| CBL | NM_007619 | 0.41 | | 0.61 | |
| CBLB | NM_001033238 | -0.47 | | -0.24 | |

| | | 12 hpi | | 24 hpi | |
|---|---|---|---|---|---|
| **Gene** | **Accession** | **ZMAD value** | **Code** | **ZMAD value** | **Code** |
| CBLC | NM_023224 | 0.00 | | -0.07 | |
| CDC42 | NM_009861 | 0.56 | | 2.42 | |
| CIB1 | NM_011870 | 1.27 | | 0.69 | |
| CIB2 | NM_019686 | 0.26 | | 0.28 | |
| CLTA | NM_001080386 | -1.08 | | -0.76 | |
| CLTB | NM_028870 | 0.56 | | -0.45 | |
| CLTC | NM_001003908 | 2.11 | | 0.77 | |
| D6ERTD32E | NM_177466 | 1.84 | | -0.09 | |
| D9BWG0185E | NM_173781 | -0.58 | | 0.00 | |
| DAB2 | NM_001008702 | -0.96 | | -1.67 | |
| DIAP1 | NM_007858 | -0.96 | | -2.16 | |
| DNM | NM_010065 | -0.16 | | -0.62 | |
| DNM2 | NM_001039520 | 0.76 | | 2.38 | |
| EEA1 | NM_001001932 | -2.26 | | -0.72 | |
| EFS | NM_010112 | 1.27 | | 0.67 | |
| EPN1 | NM_010147 | 0.28 | | 1.14 | |
| EPN2 | NM_010148 | -0.66 | | -1.33 | |
| EPN3 | NM_027984 | -0.30 | | -0.68 | |

| | | 12 hpi | | 24 hpi | |
|---|---|---|---|---|---|
| **Gene** | **Accession** | **ZMAD value** | **Code** | **ZMAD value** | **Code** |
| EPS15 | NM_007943 | -1.03 | | -2.09 | |
| EPS15-RS | NM_007944 | -1.24 | | -1.80 | |
| FYN | NM_001122892 | -0.73 | | -2.12 | |
| GORASP1 | NM_028976 | 0.15 | | -0.36 | |
| GRB2 | NM_008163 | 0.47 | | -1.19 | |
| HIP1 | NM_146001 | -0.95 | | -0.52 | |
| HIP1R | NM_145070 | 0.88 | | 0.83 | |
| Ihpk3 | NM_173027 | -0.43 | | -0.57 | |
| ITSN | NM_001110275 | 0.00 | | 0.24 | |
| LIMK1 | NM_010717 | -1.06 | | -1.70 | |
| LOC385666 | XM_001481287 | 0.61 | | 0.70 | |
| MAP4K2 | NM_009006 | 1.06 | | 0.94 | |
| MAPK8IP | NM_011162 | -0.29 | | -0.71 | |
| MAPK8IP2 | NM_021921 | 0.79 | | 0.35 | |
| MAPK8IP3 | NM_013931 | 0.60 | | 0.45 | |
| NEDD4L | NM_001114386 | 1.87 | | 0.87 | |
| NSF | NM_008740 | 0.83 | | 0.91 | |
| PACSIN1 | NM_178365 | 0.99 | | 0.24 | |
| PACSIN3 | NM_030880 | -0.19 | | 0.09 | |
| PAK1 | NM_011035 | 1.20 | | 0.47 | |

| | | 12 hpi | | 24 hpi | |
|---|---|---|---|---|---|
| **Gene** | **Accession** | **ZMAD value** | **Code** | **ZMAD value** | **Code** |
| PICALM | NM_146194 | -0.46 | | -0.14 | |
| PIK3C2G | NM_011084 | 1.20 | | 0.84 | |
| PIK3CG | NM_020272 | 0.66 | | 1.72 | |
| PIP5K1B | NM_008847 | 0.64 | | 0.45 | |
| PSCD3 | NM_011182 | 0.26 | | 0.31 | |
| RAB11A | NM_017382 | 0.26 | | -0.49 | |
| RAB11B | NM_008997 | 1.62 | | -0.67 | |
| RAB3A | NM_009001 | 1.00 | | 0.54 | |
| RAB3B | NM_023537 | 0.81 | | 0.34 | |
| RAB3C | NM_023852 | 0.69 | | 0.23 | |
| RAB3D | NM_031874 | 3.33 | | 3.13 | |
| RAB4A | NM_009003 | -0.04 | | 0.95 | |
| RAB4B | NM_029391 | 2.42 | | 1.56 | |
| RAB5A | NM_025887 | 0.84 | | 0.46 | |
| RAB5B | NM_011229 | 2.66 | | 1.21 | |
| RAB5C | NM_024456 | 2.38 | | 3.16 | |
| RAB6 | NM_024287 | 0.30 | | 0.58 | |
| RAB7L1 | NM_144875 | 0.72 | | 0.83 | |
| RAB8A | NM_023126 | 1.63 | | 0.54 | |
| RAB8B | NM_173413 | 0.61 | | 0.43 | |

| | | 12 hpi | | 24 hpi | |
|---|---|---|---|---|---|
| **Gene** | **Accession** | **ZMAD value** | **Code** | **ZMAD value** | **Code** |
| RAC1 | NM_009007 | -0.67 | | -0.45 | |
| RHOA | NM_016802 | -1.34 | | -2.60 | |
| ROCK1 | NM_009071 | 2.23 | | 0.93 | |
| ROCK2 | NM_009072 | 0.10 | | 0.15 | |
| SH3D1B | NM_011365 | 0.74 | | 0.67 | |
| SH3GLB1 | NM_019464 | 1.00 | | -0.31 | |
| SH3GLB2 | NM_139302 | -0.33 | | 0.47 | |
| SNAP91 | NM_013669 | 0.61 | | 0.26 | |
| STAU1 | NM_011490 | 0.03 | | 0.33 | |
| SYNJ2 | NM_001113351 | -0.17 | | -0.03 | |
| SYT1 | NM_009306 | 1.90 | | 2.23 | |
| SYT2 | NM_009307 | -0.23 | | -0.35 | |
| VAMP1 | NM_009496 | 0.08 | | 3.30 | |
| VAMP2 | NM_009497 | -0.34 | | -0.75 | |
| VAPA | NM_013933 | 0.88 | | 0.34 | |
| VAPB | NM_019806 | 0.21 | | 0.57 | |
| VAV2 | NM_009500 | 1.16 | | 1.88 | |
| VIL2 | NM_009510 | 0.04 | | 0.32 | |
| WAS | NM_009515 | 1.12 | | 0.32 | |
| WASF1 | NM_031877 | 1.91 | | 0.93 | |

| | | 12 hpi | | 24 hpi | |
|---|---|---|---|---|---|
| **Gene** | **Accession** | **ZMAD value** | **Code** | **ZMAD value** | **Code** |
| WASF2 | NM_153423 | -0.02 | | -0.77 | |
| WASF3 | NM_145155 | -0.01 | | 0.73 | |

**Supplemental Table 2. shRNA screen against membrane trafficking genes**

| Gene | shRNA # | P value | Gene | shRNA # | P value |
|---|---|---|---|---|---|
| Actr10 | 1 | 0.1177 | Rab27b | 1 | 0.1458 |
| Actr10 | 2 | 0.5923 | Rab27b | 2 | 0.7872 |
| Actr10 | 3 | 0.8073 | Rab27b | 3 | 0.9675 |
| Actr10 | 4 | 0.9793 | RAB27B | 4 | 0.0744 |
| Arfip2 | 1 | 0.4932 | Rab3d | 1 | 0.0819 |
| Arfip2 | 2 | 0.4412 | Rab3d | 2 | 0.3344 |
| Arfip2 | 3 | 0.7436 | Rab3d | 3 | 0.7929 |
| Arfip2 | 4 | 0.5418 | Rab3d | 4 | 0.8173 |
| Arrb1 | 1 | 0.6047 | Rab4b | 1 | 0.4803 |
| Arrb1 | 2 | 0.4322 | Rab4b | 2 | 0.3532 |
| Arrb1 | 3 | 0.8456 | Rab4b | 3 | 0.6605 |
| Arrb1 | 4 | 0.6144 | RAB4B | 4 | 0.0172 |
| CDC42 | 1 | 0.4182 | Rab5b | 1 | 0.7291 |
| CDC42 | 2 | 0.1961 | Rab5b | 2 | 0.0172 |
| Cdc42 | 3 | 0.2991 | Rab5b | 3 | 0.7451 |
| Cdc42 | 4 | 0.3578 | Rab5b | 4 | 0.5741 |
| Cltc | 1 | 0.3965 | Rab5c | 1 | 0.1399 |
| Cltc | 2 | 0.4765 | Rab5c | 2 | 0.5613 |
| Cltc | 3 | 0.532 | Rab5c | 3 | 0.4044 |
| Cltc | 4 | 0.1725 | Rab5c | 4 | 0.3601 |

| Gene | shRNA # | P value | Gene | shRNA # | P value |
|---|---|---|---|---|---|
| Dnm2 | 1 | 0.5825 | Rab8a | 1 | 0.4154 |
| Dnm2 | 2 | 0.3522 | Rab8a | 2 | 0.1749 |
| Dnm2 | 3 | 0.0558 | Rab8a | 3 | 0.5073 |
| Dnm2 | 4 | 0.2033 | Rab8a | 4 | 0.7256 |
| Exph5 | 1 | 0.9414 | Rab8b | 1 | 0.5815 |
| Exph5 | 2 | 0.6855 | Rab8b | 2 | 0.0181 |
| Exph5 | 3 | 0.8026 | Rab8b | 3 | 0.858 |
| Exph5 | 4 | 0.7788 | RAB8B | 4 | 0.4001 |
| Rab11a | 1 | 0.4995 | Rock1 | 1 | 0.4832 |
| Rab11a | 2 | 0.6372 | Rock1 | 2 | 0.006 |
| Rab11a | 3 | 0.8636 | Rock1 | 3 | 0.8686 |
| RAB11A | 4 | 0.7568 | Rock1 | 4 | 0.8534 |
| Rab11b | 1 | 0.9513 | Syt1 | 1 | 0.3277 |
| Rab11b | 2 | 0.0293 | Syt1 | 2 | 0.0346 |
| Rab11b | 3 | 0.2814 | Syt1 | 3 | 0.6141 |
| Rab11b | 4 | 0.0091 | SYT1 | 4 | 0.6419 |
| Rab11fip5 | 4 | 0.6865 | Sytl4 | 1 | 0.0549 |
| Rab11fip5 | 1 | 0.4099 | Sytl4 | 2 | 0.8449 |
| Rab11fip5 | 2 | 0.519 | Sytl4 | 3 | 0.6043 |
| RAB11FIP5 | 4 | 0.2616 | Sytl4 | 4 | 0.6921 |
| Rab27a | 1 | 0.469 | Vamp1 | 1 | 0.5823 |

| **Gene** | **shRNA #** | **P value** | **Gene** | **shRNA #** | **P value** |
|---|---|---|---|---|---|
| Rab27a | 2 | 0.0048 | Vamp1 | 2 | 0.5284 |
| Rab27a | 3 | 0.6966 | Vamp1 | 3 | 0.3935 |
| Rab27a | 4 | 0.49 | Vamp1 | 4 | 0.6897 |

**Supplemental Table3. shRNA screen against Rab11-associated genes**

| Gene | shRNA # | P value | Gene | shRNA # | P value |
|---|---|---|---|---|---|
| Exoc1 | 1 | 0.9318 | Myo5b | 1 | 0.3832 |
| Exoc1 | 2 | 0.2858 | Myo5b | 2 | 0.4357 |
| Exoc1 | 3 | 0.7806 | Myo5b | 3 | 0.6864 |
| Exoc1 | 4 | 0.0756 | Myo5b | 4 | 0.3675 |
| Exoc2 | 1 | 0.2719 | Rab11a | 1 | 0.7994 |
| Exoc2 | 2 | 0.9959 | Rab11a | 2 | 0.0745 |
| Exoc2 | 3 | 0.7262 | Rab11a | 3 | 0.4838 |
| Exoc2 | 4 | 0.8977 | RAB11A | 4 | 0.9983 |
| Exoc3 | 1 | 0.5506 | Rab11b | 1 | 0.5452 |
| Exoc3 | 2 | 0.9824 | Rab11b | 2 | 0.4713 |
| Exoc3 | 3 | 0.6841 | Rab11b | 3 | 0.0715 |
| Exoc3 | 4 | 0.4311 | Rab11b | 4 | 0.2229 |
| Exoc4 | 1 | 0.8072 | Rab11fip1 | 1 | 0.327 |
| Exoc4 | 2 | 0.4192 | Rab11fip1 | 2 | 0.6949 |
| Exoc4 | 3 | 0.0971 | Rab11fip1 | 3 | 0.0616 |
| Exoc4 | 4 | 0.8716 | Rab11fip1 | 4 | 0.0041 |
| EXOC5 | 1 | 0.5385 | RAB11FIP2 | 1 | 0.6161 |
| EXOC5 | 2 | 0.5588 | Rab11fip2 | 2 | 0.9433 |
| Exoc5 | 3 | 0.0179 | Rab11fip2 | 3 | 0.2654 |
| Exoc5 | 4 | 0.4602 | Rab11fip2 | 4 | 0.6218 |
| Exoc6 | 1 | 0.8661 | Rab11fip3 | 1 | 0.338 |

| Gene | shRNA # | P value | Gene | shRNA # | P value |
|---|---|---|---|---|---|
| Exoc6 | 2 | 0.9808 | Rab11fip3 | 2 | 0.3315 |
| Exoc6 | 3 | 0.6457 | Rab11fip3 | 3 | 0.4513 |
| Exoc6 | 4 | 0.796 | Rab11fip3 | 4 | 0.0233 |
| Exoc7 | 1 | 0.3209 | Rab11fip4 | 1 | 0.4408 |
| Exoc7 | 2 | 0.8242 | Rab11fip4 | 2 | 0.8527 |
| Exoc7 | 3 | 0.6668 | Rab11fip4 | 3 | 0.4792 |
| Exoc7 | 4 | 0.5208 | Rab11fip4 | 4 | 0.4426 |
| Exoc8 | 1 | 0.4807 | Rab11fip5 | 1 | 0.3468 |
| Exoc8 | 2 | 0.5697 | Rab11fip5 | 2 | 0.4045 |
| Exoc8 | 3 | 0.5392 | Rab11fip5 | 3 | 0.7385 |
| Exoc8 | 4 | 0.7132 | Rab11fip5 | 4 | 0.305 |
| Kif13a | 1 | 0.2845 | Sh3bp5 | 1 | 0.044 |
| Kif13a | 2 | 0.4443 | Sh3bp5 | 2 | 0.1828 |
| Kif13a | 3 | 0.7925 | Sh3bp5 | 3 | 0.7538 |
| Kif13a | 4 | 0.8098 | Sh3bp5 | 4 | 0.4745 |
| Kif5a | 1 | 0.5565 | Trappc2l | 1 | 0.3474 |
| Kif5a | 2 | 0.1294 | Trappc2l | 2 | 0.2794 |
| Kif5a | 3 | 0.9502 | Trappc2l | 3 | 0.1306 |
| KIF5A | 4 | 0.4521 | Trappc2l | 4 | 0.6209 |
| Kif5b | 1 | 0.3848 | Zfyve27 | 1 | 0.233 |
| Kif5b | 2 | 0.6111 | Zfyve27 | 2 | 0.5971 |
| Kif5b | 3 | 0.4694 | Zfyve27 | 3 | 0.7208 |

| Gene | shRNA # | P value | Gene | shRNA # | P value |
|---|---|---|---|---|---|
| Kif5b | 4 | 0.1674 | Zfyve27 | 4 | 0.0281 |
